## Supplementary material for "Changes in the abundance of Danish orchids over the past 30 years": Electronic supplements

### Appendix A: Citizen Science in Denmark

In Denmark, there is a long-time tradition for national species surveillance involving voluntary citizens. Birds were the first group of organisms to be involved in a species-monitoring program. In 1899, the Danish ornithologist Hans C.C. Mortensen (1856-1921) founded the project of ringing birds in order to follow their migration. During the 20$^{th}$ century, many Danish natural history societies launched national surveillance programs that with a modern term are called 'Citizen Science'. In 1904, the Danish Botanical Society started a complete monitoring program of the Danish flora of vascular plants – the Botanical Topographic Investigation of Denmark (TBU). The aim of the project was to gather information on the distribution and state of the Danish vascular plant flora. Officially, 262 voluntary persons participated, but more than half of them never became active (Hartvig 2015). In Denmark, 57 topographical-botanical districts were established in order to collect more detailed information on the vascular plant distribution. Field monitoring ended in 1923. The results of the monitoring were worked up and published in family order starting with the *Fabaceae* (*Papilionaceae*) in 1931 (Jessen 1931). The final scientific paper on the *Portulacaceae* and the *Valerianaceae* (Hansen and Pedersen 1976) was published in 1976. In total, 41 scientific papers have been published on the distribution of the Danish vascular plant flora. In 1989, a scientific paper (Vestergaard and Hansen 1989) on the general distribution pattern of the Danish vascular plant flora was issued, which formally closed the project.

In 1991, the Danish Botanical Society launched a follow-up project called Atlas Flora Danica (AFD) in addition to the TBU-project. The AFD project is a time-limited survey (1991-2012) of the actual flora, based on a standardized method. The survey includes all vascular plants, native as well as alien, met with outside cultivation. The recording units were 5 x 5 km squares in the UTM grid. A total of 2228 squares cover Denmark. Of the 2228 squares, 1300 were surveyed thoroughly. The project was completed in 2015 with the publication of a three-volume book containing a presentation of the recorded vascular plant species in the project and distribution maps for most of the species (Hartvig 2015).

The Buderupholm Forest District (now Himmerland Forest District) initiated the annual census of the local population of *Cypripedium calceolus* in 1943, which is one of oldest and still ongoing surveillances of a single species. The position of the clones of *C. calceolus*, which is surrounded by a fence to protect the plants from pricking of the shoots and digging of the plants, has been mapped precisely in order to follow the development of each individual clone. The number of both vegetative and flowering aerial shoots has been recorded every year since 1943, except for 1945 where only the flowering aerial shoots were counted, and 1946 where no recordings of the aerial shoots were performed at all.

Many other national terrestrial surveillance programs involving volunteer citizens have been performed in Denmark, focusing on other organism groups. Examples are: 1) the national bird-monitoring program based on Point Count Census covering the trends in the breeding population of 90 species launched in 1975 and organised by the Danish Ornithological Society. 2) monitoring of amphibians and reptiles started in 1976 and is organised by the youth society 'Natur og Ungdom'. 3) the World Wildlife Fund and the Danish Animal Welfare Society's monitoring of the Danish otter launched in 1984 and is conducted in cooperation with the Natural History Museum in Aarhus (Asbirk and Orth 1987). Besides, the Danish Ornithological Society has performed three national atlas surveys of bird species in 1971 – 1974, 1993- 1996 and the latest in 2014 – 2017 using the same 5 x 5 km squares in the UTM grid that were used in the botanical AFD-project.

### Appendix B: The National Orchid Monitoring Program

The Nature Protection Agency in cooperation with other national institutions held a symposium in October 1986 in order to get an overview of the many Danish surveillance programs run by other national agencies, county counsels, universities, and natural history societies. The aim of the symposium was to get inspiration on how to work out a national terrestrial monitoring program to support effective nature management.

At the turn of year 1986/1987, a fusion of the Forest Agency and the Nature Protection Agency resulted in the establishment of the Forest and Nature Agency under the Ministry of Environment. One of the first tasks of the new agency was to work out and launch a national monitoring program based on the recommendation of the 1986-symposium.

The National Orchid Monitoring Program, launched in cooperation with the Danish Botanical Society in 1987, is one example. The aim of the program is to monitor the state of and possible changes in orchid abundance in representative Danish orchid populations by means of annual censuses (Løjtnant 1991). Besides monitoring orchid abundance, the aim of the monitoring program is to document the effects of joint legislation on biodiversity and to collect data that may be useful in understanding the causal mechanisms underlying the observed changes in the state of the orchid populations.

The National Orchid Monitoring Program relies on volunteers to conduct the annual census of the selected orchid populations in the field. Therefore, volunteers have been recruited as surveyors together with professionals already involved in orchid censuses to carry out the annual censuses. Approximately 170 surveyors have contributed to the project over the past 30 years since its start in 1987. Some surveyors have contributed to the monitoring program from the beginning, whereas others have contributed for shorter periods or have joined the program later and are still involved. Unfortunately, a few surveyors have passed away.

In 1994, The National Environmental Research Institute (NERI) under the Ministry of Environment took over the coordination and maintenance of the monitoring program. NERI set up a database for storing the data collected in the program and ensured that data were accessible on the web (http://bios.au.dk/raadgivning/natur/planter/). When NERI was included in the Department of Bioscience at Aarhus University in 2007, the university became responsible for coordinating and maintaining the monitoring program.

### Methodology

The first coordinator of the National Orchid Monitoring Program, the Danish orchid expert Bernt Løjnant, launched four methods for performing the annual censuses of aerial shoots of flowering and vegetative orchid specimens. 1) A total count of all specimens in an orchid population. The benefit of the method is that it is able to monitor the orchid populations that may emerge at different locations from year to year. 2) A count of all specimens within a number randomly laid out, permanent squares on 1 $m^2$ or circles on 0.1 $m^2$. 3) A total count of specimens within a permanent 'strip', e.g. 50 m in length and 1 m width in an orchid population. The method has been used on a large Danish population of *Liparis loeselii* on Zealand comprising thousands of plants. 4) A total count of all specimens within a permanent plot with a size of up to 100 $m^2$. The coordinator recommended using the latter method in general (Løjtnant 1991). A disadvantage is that many orchids disappear from the permanent plots only to frequently appear in the surroundings. Besides, a permanent plot may disappear, especially plots established on plastic soil where the plot and the orchid population may possibly slide into the sea. The most applied methods in the program have proved to be methods 1 and 4, where, initially, the flowering shoots have been counted and, later, the vegetative shoots as well, when possible.

### Pressures

Another important field observation is the state of possible pressures on the orchid populations. The pressure comprises all factors, both natural and causes by human impact, that affect the growth of an orchid population at a site. Pressure is divided into five main categories. 1) Grazing. 2) Intensity of forest management. 3) Overgrowth. 4) Public disturbance. 5) Other impacts (not included in the previous four).

1) *Grazing* comprises all kinds of influence caused by grazing livestock at the orchid sites. Thus, the category does not include the influence of wild animals, e.g. red deer (*Cervus elaphus*), roe deer (*Capreolus capreolus*) and geese (*Anser anser*). Their influence is noted in category five.

2) *Forest management* is the influence caused by forestry at the orchid sites. The influence covers everything from picking of selected trees to pollarding to intensive management.

3) *Overgrowth* comprises the pressures caused by the surrounding vegetation to the growth of an orchid population at a site. Overgrowth typically occurs where the previous management has changed or has ceased. This includes all degrees of overgrowth, from increased dominance of tall-growing herbs to the growth of woody plants.

4) *Public disturbance* is the influence caused by recreational activities to orchid sites. Thus, the category comprises the degree of impact caused by such activities as random human damage to orchid populations by thread or picking flowers for a bouquet to targeted plant collection and digging of entire plants or clones for gardening or commercial use.

5) The category comprises impacts caused by the use of other management practices apart from those described in the previous categories. A typical example is hay mowing.

As mentioned in the body text, the intensity of the pressures is divided into a four-step scale defined according to the type of impact and is shown in Table B1-B4.

Unfortunately, some of the definitions are rather weakly defined, inducing a great chance for subjective interpretation by the surveyors. The bias in the interpretation has caused surveyors to doubt which category to apply and, thus, they may have chosen to report randomly between the two categories. The intermediate choice is inappropriate when the collected data should be used for analysis of the degree of pressure.

Table B1. Degrees of impact by grazing with livestock (after Wind 1999)

| **Category** | **Definition** |
|---|---|
| **None** | The site is not grazed by livestock |
| **Weak** | The vegetation cover is coherent and dominated by tall-growing herbs and woody plants. Tussock and hillock structures are only seen to a lesser extent |
| **Moderate** | The vegetation cover is coherent and dominated by low-growing, light-dependent species. Often, a tussock and hillock structure has developed with minor vegetation and less gaps |
| **Strong** | The vegetation cover has been grazed intensively and is incoherent. Because of trampling gaps in the vegetation cover have been created, tussocks and hillocks are often stepped into pieces |

Table B2. Degrees of forestry management (after Wind 1999)

| Category | Definition |
|---|---|
| **None** | The site is characterized as natural forest or untouched forest |
| **Weak** | The picking of selected trees or coppicing is carried out extensively, whereby the forest floor is only illuminated to a small extent. Forest regeneration occurs through continuous self-regeneration of trees and shrubs |
| **Moderate** | The picking of selected trees or coppicing is carried out to such a degree that the forest floor is partly illuminated. Forest regeneration occurs by strong self-regeneration of seed plants or by self-growth of trees and shrubs |
| **Hard** | Clear cutting, establishment of new cultures of woody plants, cultivation of the forest floor, use of fertilizers and pesticides and heavy pruning of the forest vegetation |

Table B3. Degrees of overgrowth (after Wind 1999)

| Category | Definition |
|---|---|
| **None** | No sign of overgrowth at the site |
| **Weak** | The overgrowth is so modest that, in a practical sense, it does not affect the growth of the orchid population |
| **Moderate** | The overgrowth is such that it will affect the growth of the orchid population in the long term |
| **Hard/strong** | The overgrowth is such that it will affect the growth of the orchid population in the short term |

Table B4. Degrees of overgrowth (after Wind 1999)

| Category | Definition |
|---|---|
| **None** | Unintended traffic, pricking or digging of orchids have not been recorded |
| **Weak** | Unintended traffic in an orchid population has been recorded |
| **Moderate** | Unintended traffic, pricking or digging of orchids have been recorded to a moderate extent |
| **Hard/strong** | Targeted digging of parts of or entire populations of orchids or a great degree of traffic have been recorded |

Finally, the surveyor must assess the suitability of the pressure on the orchid population. The aim of the assessment of the suitability is to see weather the intensity of the pressure is good for or will harm the growth of the orchid population at the site in question. Many orchid species require balanced regulation of vegetation height and extent at the site in order for the population to thrive. For example, a certain grazing pressure may cause the low-growing vegetation at a given site to be maintained when the combination of bite and tramp is optimal. If grazing pressure is too high, it may cause damage to the vegetation cover in the form of abrasion and curbing, while too low grazing pressure may cause problems in the form of overgrazing.

The suitability of the intensity of the impact depends on the ecology of each orchid species. Some species thrive best in direct sunlight, while others prefer shade. For example, the number of *Orchis mascula* and *Neottia ovata* increases by the time of application, while *Epipactis* and *Cephalanthera* species thrive in untouched forest (Wind 1989).

When the surveyors have finished the field work, all information is filled in data sheets that are sent to the program manager at Aarhus University no later than at the end of the season. The program manager examines the received sheets, making a quality check of the information to secure its validity prior to the data being stored in the orchid database.

### Appendix C: Statistical model

The two models were implemented in R-INLA as

Model1=Present~Year+f(Site,model="iid") (S1),

Model2=Present~Year+f(Site,model="iid")+f(Site1,Year,model="iid") (S2).

Both model were fitted by, fit=inla(Model, data, family="poisson", E=1, control.compute=list(dic=TRUE)) (S3).

Asbirk S, Orth H. 1987. Naturovervågning – rapport fra et symposium i middelfart. Hørsholm: Skov- og Naturstyrelsen.

Hansen A, Pedersen A. 1976. Portulacaceernes og valerianaceernes udbredelse i danmark. Bot Tidsskr. 71:57-74.

Hartvig P. 2015. Atlas flora danica. København: Gyldendal.

Jessen K. 1931. The distribution within denmark of the higher plants. Ii. The distribution of the papilionaceae within denmark. D Kgl Danske Vidensk Selsk Skrifter Naturvidensk Og Mathm. 9.

Løjtnant B. 1991. Overvågning af orchideer 1987-89. Flora og Fauna. 97:63-121.

Vestergaard P, Hansen K. 1989. Distribution of vascular plants in denmark. Opera Botanica. 96:1-163.
